# Dose-dependent effects of histone methyltransferase NSD2 on site-specific double-strand break repair

**DOI:** 10.1101/2023.10.18.562991

**Authors:** Koh Iwasaki, Akari Tojo, Haruka Kobayashi, Yoshitaka Kamimura, Yasunori Horikoshi, Atsuhiko Fukuto, Jiying Sun, Manabu Yasui, Masamitsu Honma, Atsushi Okabe, Ryoji Fujiki, Atsushi Kaneda, Satoshi Tashiro, Akira Sassa, Kiyoe Ura

## Abstract

Histone modifications are catalyzed and recognized by specific proteins to regulate dynamic DNA metabolism processes. NSD2 is a histone H3 lysine 36 (H3K36)-specific methyltransferase that associates both with various transcription regulators and DNA repair factors. Specifically, it has been implicated in the repair of DNA double-strand breaks (DSBs); however, the role of NSD2 during DSB repair remains enigmatic. Here, we show that NSD2 does not accumulate at DSB sites and that the localization of NSD2 at chromatin is maintained even after DSB formation. Using three different DSB repair reporter systems, which contained the endonuclease site in the active thymidine kinase gene (*TK*) locus, we demonstrated separate dose-dependent effects of NSD2 on HR, canonical-NHEJ (c-NHEJ), and non-canonical-NHEJ (non-c-NHEJ). Endogenous NSD2 has a role in repressing non-c-NHEJ, without affecting DSB repair efficiency by HR or total NHEJ. Furthermore, overexpression of NSD2 promotes c-NHEJ repair suppressing HR repair. Therefore, we propose that NSD2 has functions in chromatin integrity at the active regions during DSB repair.

## 1 Introduction

DNA double-strand breaks (DSBs) are considered very dangerous lesions since DSBs have the potential to induce harmful genetic alterations, such as local DNA insertions/deletions or chromosomal rearrangements. To maintain genomic integrity, cells respond to DSBs by assembling various factors that repair this damage by either homologous recombination (HR) or non-homologous end joining (NHEJ) (Ceccaldi et al., 2016; Chiruvella et al., 2013; Clouaire and Legube, 2019). Moreover, NHEJ consists of two pathways, canonical-NHEJ (c-NHEJ) pathway and non-canonical-NHEJ (non-c-NHEJ). During DSB repair, broken DNA ends are protected from resection and directly ligated in the c-NHEJ pathway. In contrast, in the non-c NHEJ pathway, which includes microhomology-mediated end-joining (MMEJ), alternative end-joining (a-EJ), and single-strand annealing (SSA), end resection is required to generate single-stranded DNA tails, thereby making it more error-prone than c-NHEJ (Chiruvella et al., 2013; Zhao et al., 2020). The packaging of eukaryotic DNA into chromatin affects all dynamic processes of DNA metabolism. The assembly of nucleosomes—the basic subunit of chromatin structure—restricts the access of DNA binding factors to their target sites (Luger et al., 1997; Wolffe, 2000), and modulates chromosomal environments through chromatin modifications (Felsenfeld and Groudine, 2003; Allis et al., 2015). The diverse methylation of histone lysine residues is recognized and catalyzed by specific proteins that play critical roles in the regulation of genetic information (Allis et al., 2015; Janssen and Lorincz, 2021).

NSD2 (also known as WHSC1 or MMSET) is a mammalian histone H3 lysine 36 (H3K36)-specific histone methyltransferase (HMT), which has been considered as both an H3K36 mono-and dimethyl (1/2)-specific HMT, despite its H3K36me3 activity (Kuo et al., 2011; Lam et al., 2022; Nimura et al., 2009). Defects in *NSD2* are linked to Wolf– Hirschhorn syndrome, in which patients show various developmental defects, including growth delays, whereas the overexpression of *NSD2* is detected in multiple myelomas and many solid tumors (Bennett et al., 2017; Li et al., 2019). The link between *NSD2* and the pathogenesis of human diseases indicates the importance of the NSD2 cellular functions in mammals. NSD2 collaborates with transcription factors and chromatin regulators to tune gene expression (Nimura et al., 2009; Ouda et al., 2018; Sarai et al., 2013). Homozygous *Nsd2*-deficient mice show abnormal gene regulation and die early after birth (Campos-Sanches, et al., 2017; Dobenecker et al., 2020; Nimura et al., 2009; Zhang et al., 2022). Furthermore, NSD2 has been implicated in the repair of DSBs. Enhanced DSB damage repair was observed by pathological NSD2 overexpression in multiple myeloma cells (Shah et al., 2016). Although NSD2 was first reported to recruit 53BP1, mediated by H4K20 methylation, to promote NHEJ, this model has been subsequently argued for by enzymatic and genetical studies (Hartlerode et al., 2012; Panier and Boulton, 2014; Pei et al., 2011). The accumulation of NSD2 at DSBs was also shown to promote the recruitment of RAD51, suggesting NSD2 is involved in DSB repair through the HR pathway (Wang and Goldstein, 2016). Conversely, NSD2 promotes ligase 4-dependent c-NHEJ at uncapped telomeres through unknown mechanisms (Krijer et al., 2020). Accumulating data suggest a link between NSD2 and DSB repair, although the function of NSD2 in the repair of DNA damage remains enigmatic.

In this study, we aimed to explore the role of NSD2 in DSB repair. Firstly, we demonstrated that the localization of NSD2 at chromatin was maintained even after DNA damage formation. NSD2 catalyzes H3K36me associated with the transcriptionally active state (Lam et al., 2022). Since it localizes at euchromatin, excluding DAPI-stained heterochromatic foci in the mouse ES cells (Nimura et al., 2009), we investigated the effects of NSD2 on DSB repair at the constitutively active thymidine kinase gene (*TK*) locus in human B lymphoblastoid TK6 cells. The three TK6-derived cell lines contain different endonuclease I-*Sce*I sites and mutations in the endogenous *TK* gene; thus, the positive/negative drug selection assays for the TK phenotypes enabled the separate DSB repair pathways to be traced separately (Honma et al., 2003, 2007). We established *NSD2*^-/-^ and clones overexpressing NSD2 from each TK6-derived cell line. The *NSD2^-/-^*clones exhibited activation of the non-c-NHEJ without inducing significant changes in the efficiency of HR or total NHEJ; moreover, the overexpression of NSD2 facilitated DSB repair by NHEJ and suppressed HR. Furthermore, we found that the number of ionizing radiation (IR)-induced RAD51 foci was reduced depending on the NSD2 expression levels. Our results suggest that NSD2 represses end-resection at the DSBs and large excess amounts of NSD2 promote c-NHEJ at transcriptionally active regions. We propose a novel role for NSD2 in the maintenance of chromatin during DSB repair.

## 2 Results

### 2.1 The localization of NSD2 at chromatin was maintained even after DNA damage formation

Since several groups have reported that NSD2 accumulates at DSBs, where it recruits repair factors, such as 53BP1 and RAD51 (Pei et al., 2011; Wang and Goldstein, 2016), we first examined the dynamics of NSD2 after DSB formation in human fibroblast cells. Endogenous NSD2 localized to the nucleoplasm without forming specific foci. Although 53BP1, which localized mainly to the nuclear bodies besides the nucleoplasm before irradiation, accumulated at the damaged sites 30 min after microirradiation, we could not observe significant changes of the localization of NSD2 even 2 hours after micro-irradiation (Figure 1a). We further examined the dynamics of NSD2 following DNA damage by investigating the fluorescence recovery of EGFP-NSD2 in two independent strips of a single nucleus immediately after micro-irradiation: one in the irradiated area and the other in an unirradiated region (Figure 1b). No substantial difference was observed in the fluorescence recovery speed between the two regions in the cell. The fluorescence intensities in both the irradiated and unirradiated regions were almost recovered within 2 min after bleaching. NSD2 contains various chromatin binding domains, including two PWWP domains, and stably associates with chromatin (Nimura et al., 2009, Sankaran et al., 2016). Our findings suggest that the mobility of NSD2 is not affected by DNA damage formation and a stable association between NSD2 and chromatin is maintained during DSB repair.

**Figure 1.**
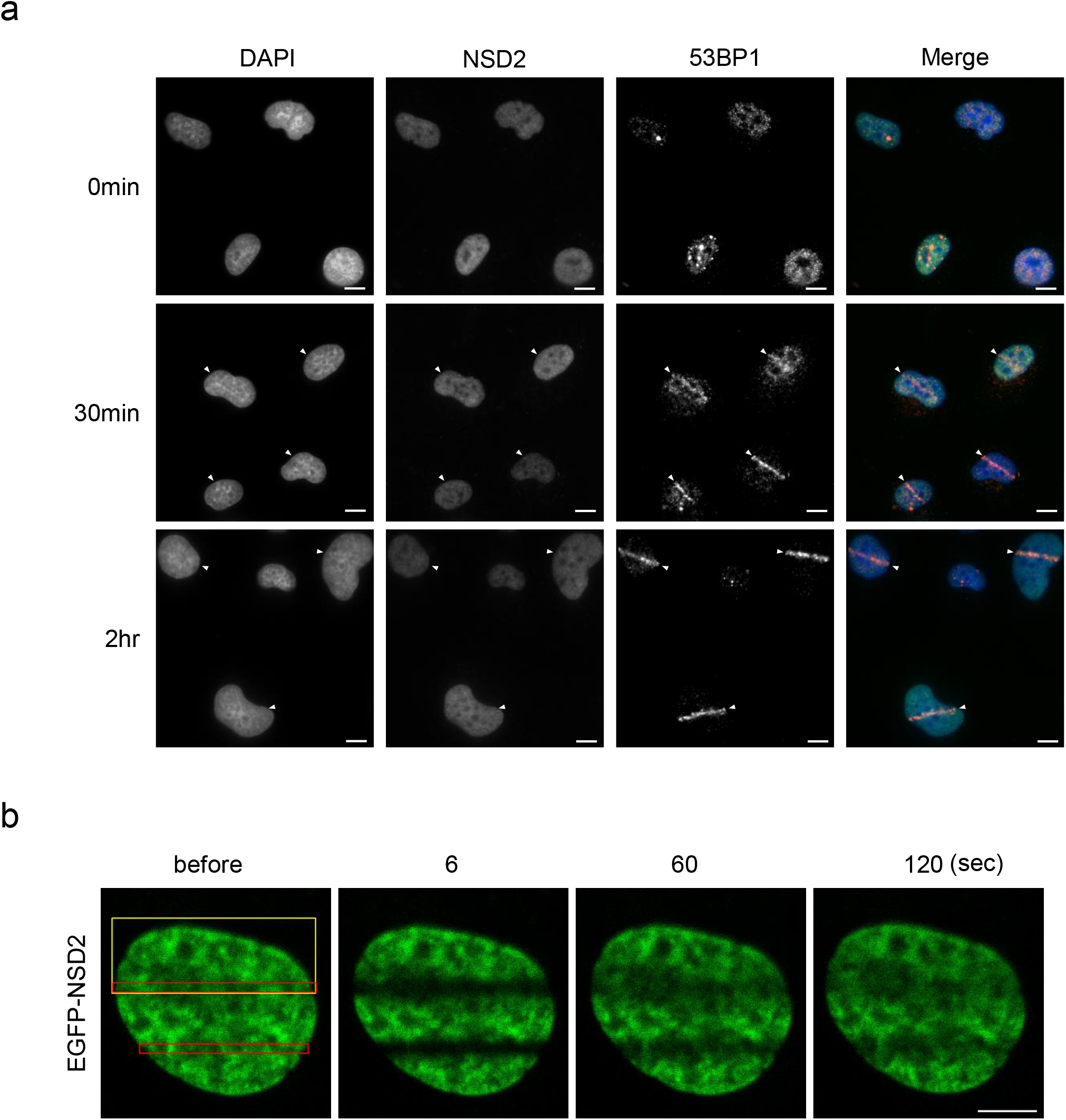
NSD2 does not accumulate at sites of DSB. (a) Immunofluorescence staining of GM0637 cells at 30 minutes and 2 hours after UVA micro-irradiation, using anti-NSD2 and anti-53BP1 antibodies. NSD2, 53BP1, and DNA (DAPI) are shown in green, red, and blue, respectively, and merged. Arrowheads indicate the micro-irradiation sites. Scale bars: 10 μm. (b) FRAP analysis to monitor NSD2 dynamics at damage sites. EGFP-NSD2-expressing GM10637 cells were first micro-irradiated (yellow boxes), and then, photobleached (red boxes). The EGFP signal was examined before and at the indicated times after micro-irradiation. Scale bars: 5 μm.

### 2.2 *NSD2*-deficiency reduces cell growth and H3K36me2 and H3K36me3 methylation levels

To analyze the role of NSD2 in each DSB-repair pathway, we first disrupted *NSD2* before stably overexpressing it in three different DSB repair reporter TK6-derived cell lines TSCER2, TSCE105, and TSCE5 (Figures 3a, 3c, and 4a).

Three *NSD2^-/-^* cell lines grew significantly more slowly than wildtype cells. The increased doubling time of *NSD2-*deficient cells was recovered when NSD2 was stably overexpressed in these cells (Figure 2a, 2c, and 2e). We found that the levels of both H3K36me2 and H3K36me3 were reduced in *NSD2^-/-^*cells compared to wildtype cells (Figure 2b, 2d, and 2f). In contrast, the H3K36me2/3 levels were clearly increased in the cells overexpressing NSD2. Consistent with previous reports, overexpression of NSD2 reduced the level of H3K27me3, yet did not significantly alter H4K20me2 (Figure 2b). These data indicate that NSD2 regulates H3K36me2 and H3K36me3 in TK6 cells.

**Figure 2.**
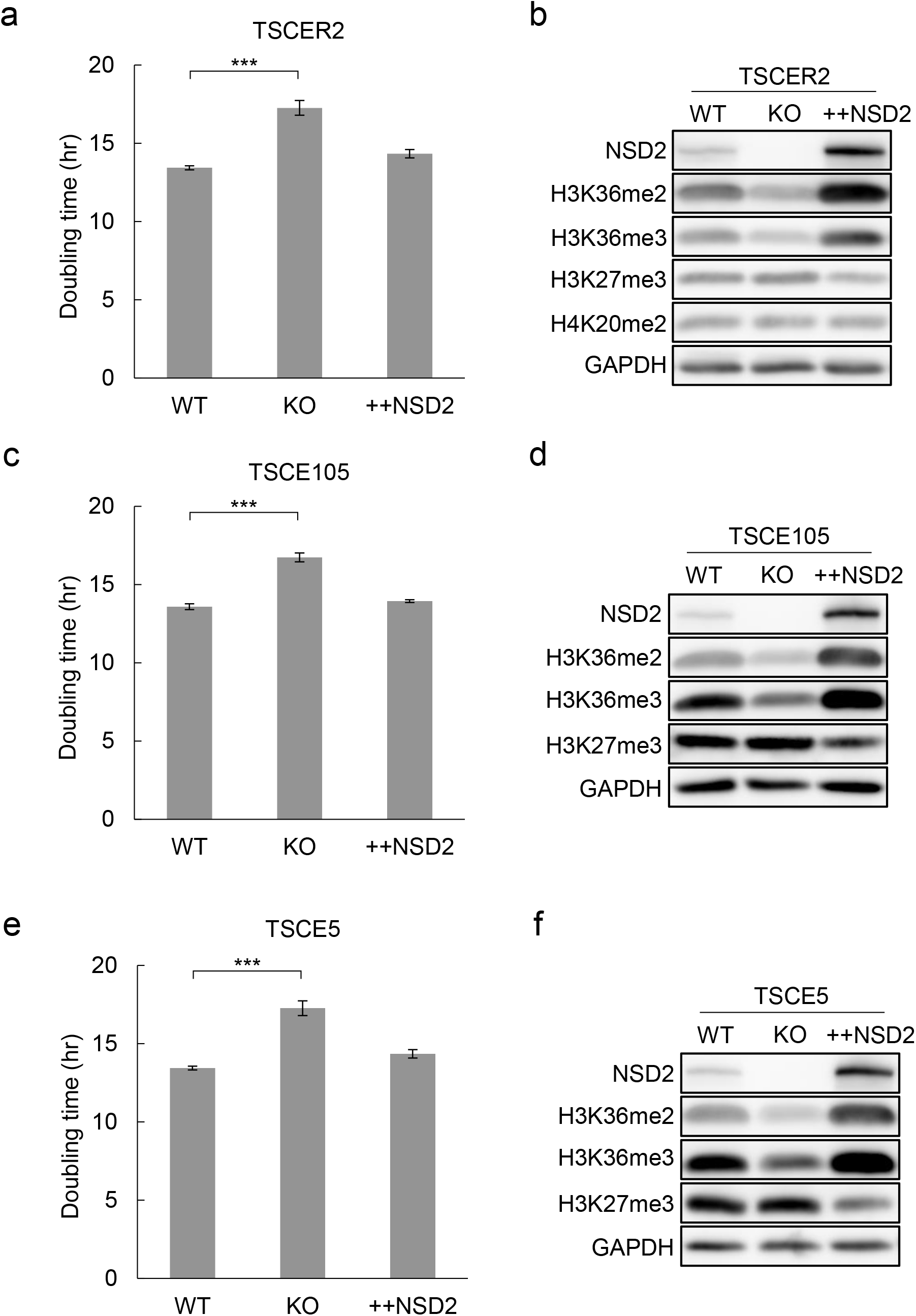
*NSD2*-deficient cells slow down cell growth and reduce the methylation levels of H3K36me2/3. (a) (c) (e) Doubling time of *NSD2*-deficient (KO) and *NSD2*-overexpressing (++NSD2) cells compared to wildtype cells (WT). Three TK6-drived cell lines, TSCER2, TSCE105, and TSCE5 are indicated. Error bars represent S.D. from n = 3 (TSCER2, TSCE105), n = 5 (TSCE5). Dunnett’s test was performed to compare all samples with WT cells. \*\*\**P* < 0.001. (b) (d) (f) The expression levels of NSD2 and the methylation status of histone H3K36me in wildtype (WT), *NSD2*-deficient (KO), and *NSD2*-overexpressing (++NSD2) cells. Three TK6-drived cell lines, TSCER2, TSCE105, and TSCE5 are indicated. Whole-cell extracts were analyzed by immunoblotting and the blots were probed with the indicated antibodies.

**Figure 3.**
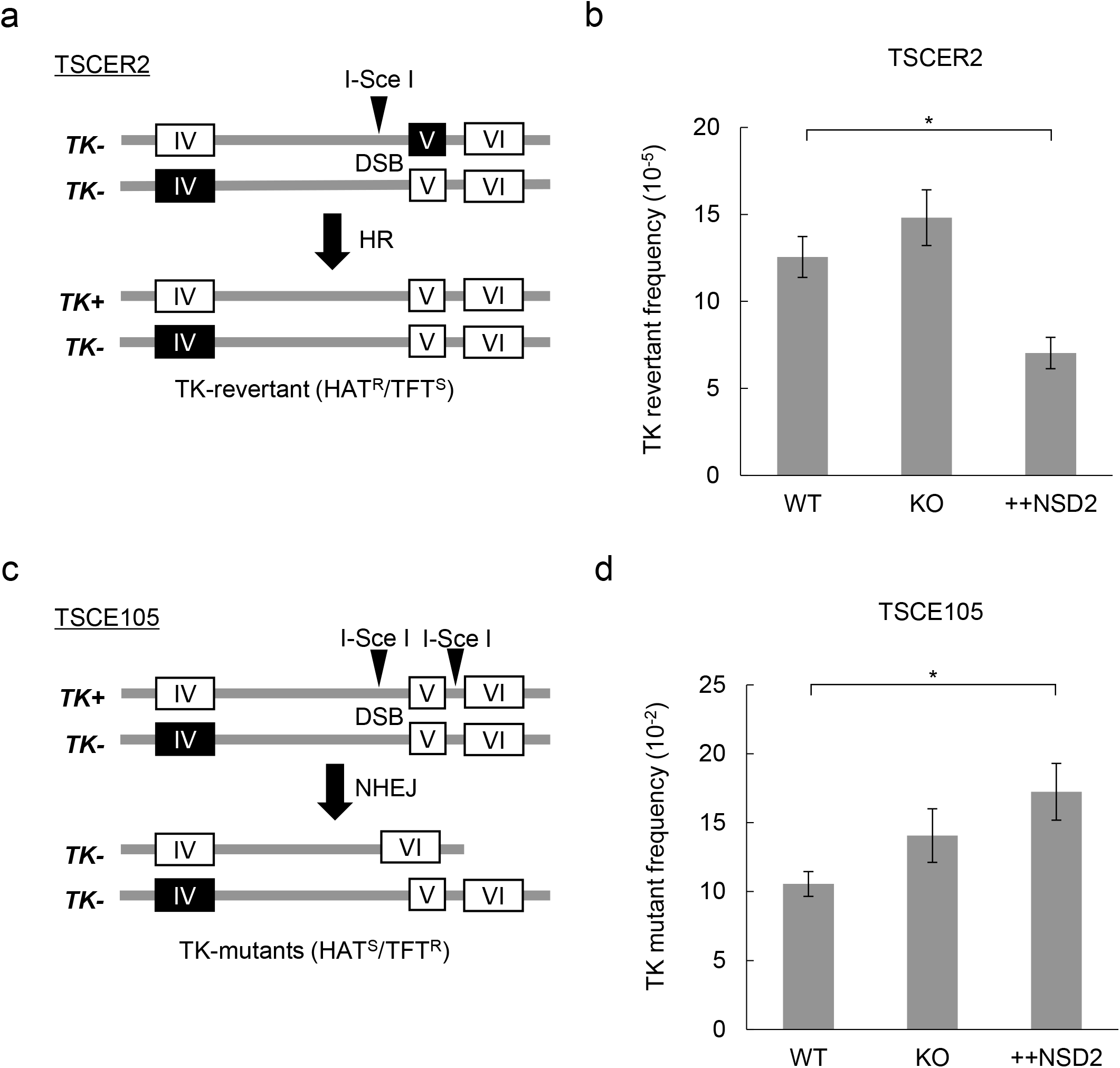
*NSD2*^-/-^ cells show no significant defects in DSB repair but overexpression of NSD2 promotes NHEJ and suppresses HR. (a) Schematic representation of the homologous recombination (HR) assay system at the *TK* locus in TSCER2 cells. Opened and closed rectangles represent the wildtype and mutant exons of TK gene, respectively. *TK*^-/-^ TSCER2 cells possess an I-*Sce*I site in intron 4 (IV). (b) Frequency of revertant mutations by HR at *TK* locus, as shown in (A), in wildtype (WT), *NSD2*-deficient (KO), and *NSD2*-overexpressing (++NSD2) cells. TSCER2-TK6 cells. Error bars represent S.D. from n = 9 for WT, n = 7 for KO, and n = 4 for KO+NSD2 cells. Dunnett’s test was performed to compare all samples with WT cells. \**P* < 0.05. (c) Schematic representation of the non-homologous end joining (NHEJ) assay system at the *TK* locus in TSCE105 cells. Opened and closed rectangles represent the wildtype and mutant exons of the TK gene, respectively. *TK*^+/-^ TSCE105 cells possess two I-*Sce*I sites that flank exon 5 in the functional allele. Both canonical-NHEJ and noncanonical-NHEJ for I-*Sce*I-induced DSBs would yield *TK*^-/-^ clones from *TK*^+/-^ cells. The number of *TK*^-/-^ clones was measured using the number of trifluorothymidine (TFT)-resistant colonies. (d) Frequency of DSB-repair mutation at *TK* locus by NHEJ in wildtype (WT), *NSD2*-deficient (KO), and *NSD2*-overexpressing (++NSD2) TSCE105-TK6 cells. Error bars represent S.D. from n = 9 for WT, n = 7 for KO, and n = 4 for KO+NSD2 cells. Dunnett’s test was performed to compare all samples to WT cells. \**P* < 0.05.

### 2.3 *NSD2*-deficiency does not reduce the efficiency of HR at the *TK* locus

We investigated the effects of NSD2 on HR at the *TK* gene locus in TSCER2 cells. The TK6-derived TSCER2 cell line is compound heterozygous (*TK*^-/-^) and contains a single I-*Sce*I endonuclease site in intron 4 of the *TK* gene (Figure 3a). Only when I-*Sce*I-induced DSB is repaired by the HR pathway, TK-proficient revertants (*TK*^+/-^) are generated and selected in the HAT medium. The revertant frequency is detectable by counting drug-resistant colonies.

We introduced the I-*Sce*I expression vector into the TSCER2 cells to generate a single DSB in the *TK* gene. In this HR reporter system, a reduction in the HR efficiency decreases the TK revertant frequency. We found that a deficiency in *NSD2* did not cause a reduction but rather a slight increase in TK revertant frequency. In contrast, NSD2 overexpression significantly reduced the number of HAT-resistant TK revertant colonies (Figure 3b). These results show that endogenous NSD2 did not induce any significant effects on HR efficiency at the *TK* gene, although it was previously reported following analysis using a DR-GFP/U2OS HR reporter assay that downregulation of NSD2 decreased HR efficiency (Wang and Goldstein, 2016). Conversely, our data demonstrated that overexpression of NSD2 suppressed HR DSB repair in an active *TK* gene.

### 2.4 NSD2 overexpression facilitates DSB repair by NHEJ

Next, we investigated the effects of NSD2 on the efficiency of NHEJ-mediated repair in the TSCE105 cell line, which is heterozygous (*TK* ^+/-^) and contains two I-*Sce*I recognition sites that tandemly flank exon 5 (Figure 3c). When DSBs that were induced by I-*Sce*I are repaired by c-NHEJ or non-c-NHEJ, exon 5 is deleted to generate *TK*-deficient mutants (*TK*^-/-^) that form colonies in the negative selection medium containing trifluorothymidine (TFT) (Honma et al., 2007).

After I-*Sce*I-induced DSB formations, we compared the NHEJ efficiency among three TSCE105 cell lines that differentially express NSD2. As seen in the HR-reporter TSCER2 cell lines (Figure 3b), we did not observe any significant differences in the *TK* mutant frequency between the wildtype and *NSD2*-deficient TSCE105 cells, indicating that endogenous NSD2 had no significant effect on the NHEJ efficiency at the *TK* gene (Figure 3d). However, interestingly, we found that overexpression of NSD2 increased the *TK* mutant frequency (Figure 3d). It should be noted that the *TK* mutant frequency in wildtype TSCE105 cells ((1.1 ± 0.090) × 10^-1^) was almost 1000 times higher than the revertant frequency in wildtype HR-reporter TSCER2 cells ((1.3± 0.12) × 10^-4^), reflecting the balance of DSB repair pathway choice in human cells. Collectively, these data indicated that DSBs were mainly repaired via NHEJ in the active *TK* gene, which was further enhanced by the large excess amount of NSD2.

### 2.5 *NSD2*-deficient cells promote non-c-NHEJ-mediated DSB repair

To further investigate the effects of NSD2 on NHEJ, which is associated with large DNA deletion, we analyzed the efficiency of non-canonical NHEJ in TSCE5 cells. The TK6-derived TSCE5 cell line is heterozygous (*TK* ^+/-^) and contains a single I-*Sce*I endonuclease site in intron 4 of the *TK* gene (Figure 4a). When DSBs induced by I-*Sce*I are repaired by non-c-NHEJ, where there is a deletion of more than 90 bp from the I-*Sce*I site in TSCE5 cells, exon 5 is inactivated to generate *TK*-deficient mutants (*TK*^-/-^) that form colonies in the negative selection medium containing trifluorothymidine (TFT).

**Figure 4.**
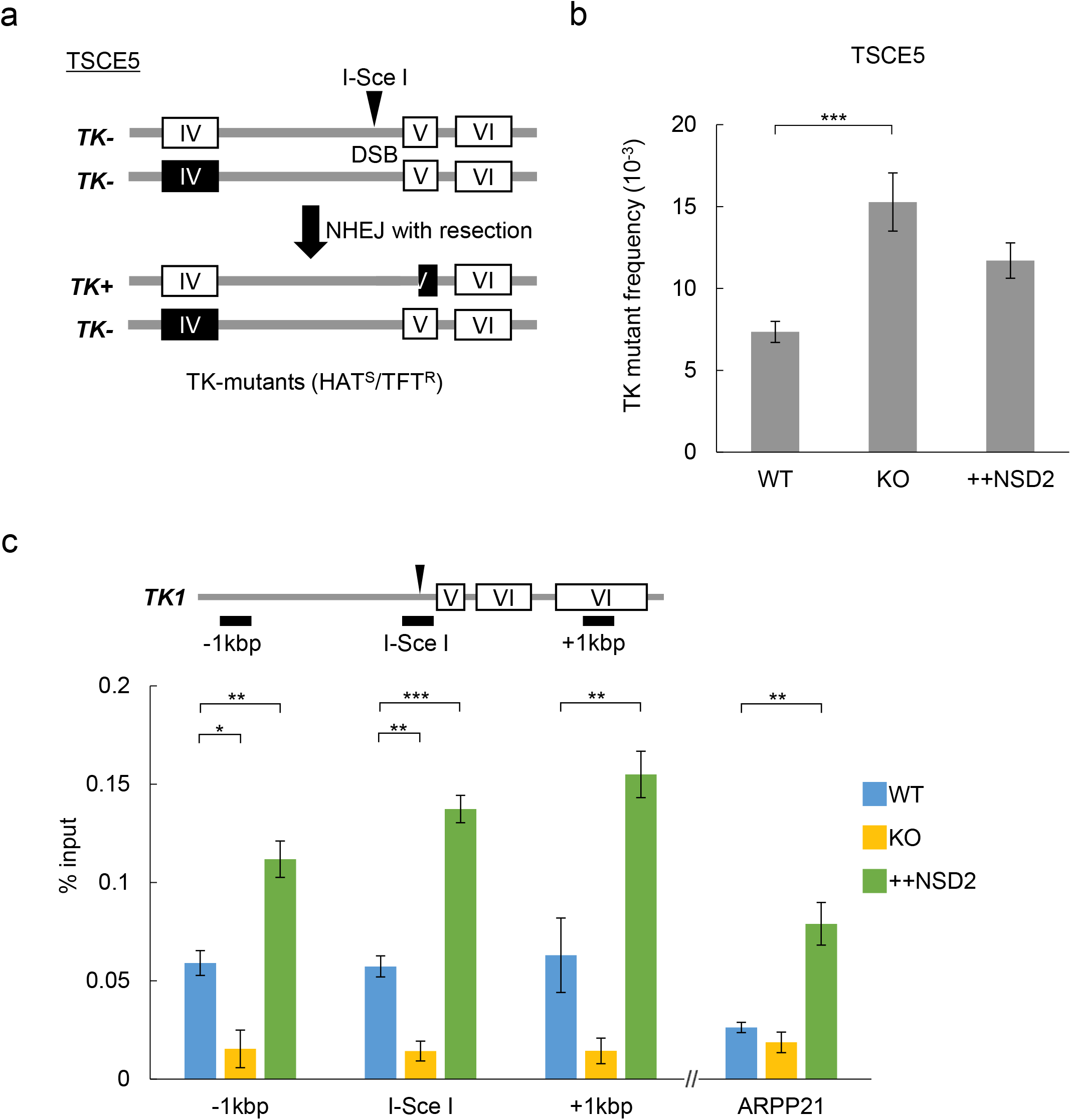
NSD2 represses non-c-NHEJ repair at the DSB site. (a) Schematic representation of the non-c-NHEJ assay system at the *TK* locus of TSCE5 cells. Opened and closed rectangles represent the wildtype and mutant exons of TK gene, respectively. *TK*^+/-^ TSCE105 cells possess two I-*Sce*I sites that flank exon 5 in the functional allele. Non-canonical NHEJ for I-*Sce*I-induced DSBs would yield *TK*^-/-^ clones from *TK*^+/-^ cells. The number of *TK*^-/-^ clones was measured by the number of trifluorothymidine (TFT)-resistant colonies. (b) Frequency of DSB repair mutations at *TK* locus by non-canonical NHEJ in wildtype (WT), *NSD2*-deficient (KO), and NSD2-rescued *NSD2*-deficient (KO+NSD2) TSCE5-TK6 cells. Error bars represent S.D. from n = 10 for WT, n = 8 for KO, and n = 3 for KO+NSD2. Dunnett’s test was performed to compare all samples with WT cells. \*\*\**P* < 0.001. (c) NSD2 occupancy around the I-*Sce*I site on the *TK* gene and silent *ARPP21* gene in wildtype (WT), NSD2-/-(KO), and *NSD2*-overexpressing (++NSD2) cells was analyzed by ChIP assays. Black rectangles indicate locations of genomic regions analyzed by ChIP assay at the *TK* gene. Error bars represent S.D. from n = 3. Dunnett’s test was performed to compare all samples with WT cells. \**P* < 0.05. \*\**P* < 0.01. \*\*\**P* < 0.001.

We found that the number of *TK*^-/-^ clones in *NSD2*-deficient cells after DSB repair was increased by almost two-fold compared to the wildtype TSCE5 cells (Figure 4b). Moreover, *NSD2* overexpression rescued the increasement in TK-mutant frequency in *NSD*2^-/-^ cells. These results suggest that NSD2 localized around the I-*Sce*I site in the *TK* gene and inhibited the non-canonical NHEJ pathway in wildtype cells during I-*Sce*I-induced DSB repair.

Consistent with this idea, sequence analysis of the I-*Sce*I site after DSB induction showed that a deficiency in NSD2 tended to enhance the DNA deletion rate. In contrast, the overexpression of NSD2 inhibited deletion around the DSB sites (Figure S1). Furthermore, we performed ChIP assays, which found that the endogenous NSD2 protein was enriched around the I-*Sce*I site in the active *TK* gene of wildtype cells, compared to the silent *ARPP21* locus (Figure 4c). The accumulation of NSD2 was reduced around the I-*Sce*I site in *NSD2*-deficient cells, although not at the *ARPP21* locus. In contrast, NSD2-binding was drastically increased in cells overexpressing NSD2 at both the *TK* locus and the *ARPP21* locus.

Overall, we concluded that NSD2-binding represses DSB repair by end-resection associated with non-c-NHEJ at the *TK* locus.

### 2.6 NSD2 repressed RAD51 foci formation after ionizing radiation exposure

As shown in Figure 1a, NSD2 is widely distributed in the nucleus. Based on our I-*Sce*I-induced mutation analyses using three different TK6-derived cell lines, we hypothesized that NSD2 more generally protects the ends of the DNA ends and suppresses HR at the sites of DSBs. Thus, to examine this possibility, we irradiated TK6 cells and investigated the effects of NSD2 on RAD51 foci formation: a marker that is mainly involved in the HR repair pathway for DSBs. RAD51 has been shown to bind to single-stranded DNA (ssDNA) and can interact with the ssDNA-binding protein RPA and RAD52 (Shima et al., 2013).

There was a limited number of RAD51 and γH2AX foci, a known DNA damage marker, in the nuclei of all three TSCE5 cell lines prior to exposure to ionizing radiation (IR) and irrespective of NSD2 expression levels (Figure 5a). However, after an hour of X-ray irradiation, the number of foci had drastically increased, even in *NSD2*^-/-^ cells (Figure 5b), which was interesting since it had been previously reported that NSD2 was required for DSB-dependent RAD51 foci formation (Wang and Goldstein, 2016). Here, we found that the number of IR-induced RAD51 foci in *NSD2*-deficient cells was slightly, yet significantly higher than in the parental wildtype cells. In contrast, the number of RAD51 foci in the cells overexpressing *NSD2* was significantly lower than in the wildtype cells (Figure 5c). Notably, there were no significant differences in the observed IR-induced γH2AX foci numbers among the three TSCE5 cell lines differentially expressing NSD2 (Figure 5d). These results demonstrate that NSD2 can suppress RAD51 foci formation after IR without affecting the global DSB repair efficiency, thereby supporting our hypothesis that NSD2 inhibits the end-resection of DSBs to suppress non-c-NHEJ at its binding sites.

**Figure 5.**
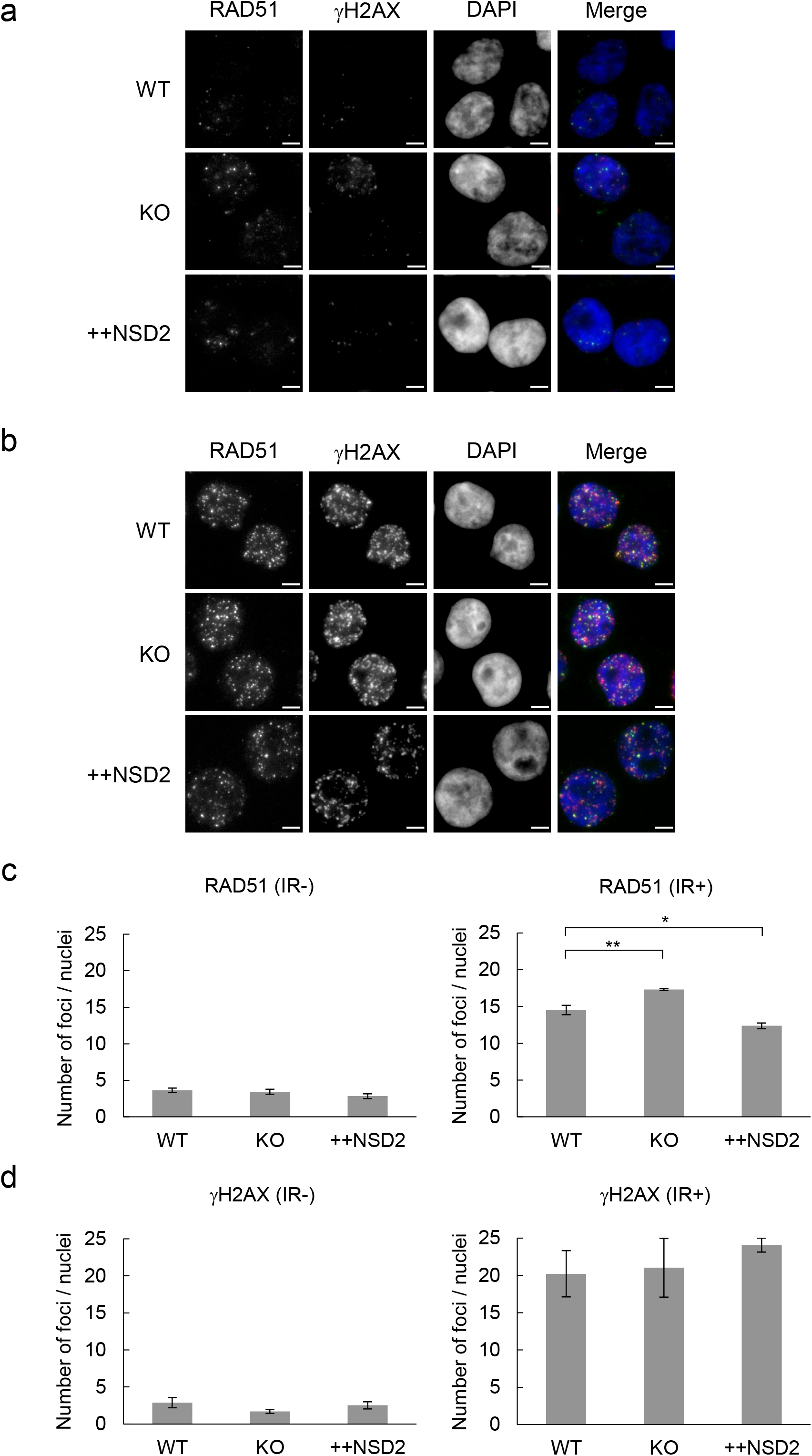
IR-induced RAD51 foci formation is enhanced in *NSD2*-deficient cells. (a) (b) Representative images of immunofluorescence analysis of wildtype (WT), *NSD2*^-/-^ (KO), and NSD2-overexpressing (++NSD2) TSCE5 cells using anti-RAD51 and anti-γH2AX antibodies before IR and one hour after IR (5 Gy). RAD51, γH2AX, and DNA (DAPI) are shown in green, red, and blue, respectively, and merged. Scale bars: 5 μm. (c) Average number of γH2AX-positive foci per nucleus before and after IR in three indicated TSCE5 cell lines. (d) Average number of RAD51-positive foci per nucleus before and after IR in three indicated TSCE5 cell lines (at least 1000 nuclei each). Dunnett’s test was performed to compare all samples with WT cells. \**P* < 0.05, \*\**P* < 0.01.

## 3 Discussion

In this study, we revealed that NSD2 is involved in the regulation of DSB repair, thereby reflecting its expression levels. Endogenous NSD2 has a role in repressing non-c-NHEJ, without affecting the HR DSB repair efficiency or total NHEJ at the active gene locus. Furthermore, NSD2 overexpression promoted the NHEJ repair pathway and suppressed the repair by HR. Although NSD2 has been implicated in DSB repair and various roles have been reported during DNA repair (Hajdu et al., 2011; Pei et al., 2011, Wang and Goldstein, 2016), how NSD2 affects each DSB pathway has not been well established, while the function of NSD2 in DNA repair is unclear. Using three different reporter systems of DSB repair, which integrated the endonuclease I-*Sce*I site in the active *TK* gene locus, we successfully demonstrated how NSD2 separately affects the HR, NHEJ, and non-c-NHEJ repair pathways. Our current study suggests different functions of NSD2 depending on the dose, which can explain how cancer cells overexpressing NSD2 can be resistant to DNA-damaging drugs and indicates the biological significance of a proper amount of NSD2 to maintain genome integrity.

NSD2 was reported to accumulate at DSBs and recruit 53BP1 or RAD51 through H4K20-dimethylation (Pei et al., 2011; Wang and Goldstein, 2016). However, we did not observe any dynamic movements of either endogenous NSD2 or EGFP-NSD2 to the DSBs after micro-irradiation, despite the accumulation of 53BP1 to DSBs (Figure 1).

Recombinant NSD2 protein catalyzes H3K36me in vitro but not H4K20me (Li et al., 2009; Nimura et al., 2009). Recent structural studies on NSD2 bound to nucleosomes have also demonstrated H3K36-specific enzymatic activity (Li et al., 2021; Sato et al., 2021). Furthermore, the accumulation of 53BP1 to DSBs or uncapped telomeres was not affected by NSD2 depletion (Hartlerode et al., 2012; Krijger et al., 2020). NSD2 contains two PWWP domains and stably associates with chromatin (Sankaran et al., 2016). Therefore, the stable association of NSD2 to chromatin seems to be maintained even after DNA damage formation, meaning NSD2 could function around the original region during DSB repair.

Since NSD2 localizes to euchromatin, excluding DAPI-stained heterochromatic foci (Nimura et al., 2009), we examined the effects of NSD2 on DSB repair at the actively transcribed *TK* gene locus, where NSD2 binds to chromatin (Figure 4c). Although NSD2 was previously reported to promote HR repair by inducing RAD51 foci formation (Wang and Goldstein, 2016), we did not observe any significant effects by endogenous NSD2 on the efficiency of either HR (Figure 3b) or NHEJ repair (Figure 3d). However, perhaps interestingly, we found that *NSD2^-/-^* cells promoted the repair of DSBs by non-c-NHEJ, which is associated with DNA end-resection (Figure 4b). Indeed, NSD2-deficient cells formed DSB-dependent RAD51 foci more than in the wildtype cells (Figure 5) and DNA deletion was enhanced around the DSB sites (Figure S1). NHEJ is the predominant repair mechanism of DSB throughout the cell cycle in mammals and can be roughly divided into major c-NHEJ and minor non-c-NHEJ. The latter, non-c-NHEJ, resects the DSB ends to generate 3’ single-stranded DNA followed by annealing to repair, thereby making it more error-prone than c-NHEJ (Chiruvella et al., 2013: Ramsden et al., 2022). Although NSD2 is not required for the recruitment of 53BP1 to the site of DSBs (Hartlerode et al., 2012; Krijger et al., 2020), endogenous NSD2 might repress the end resection of DSBs at transcriptionally active regions (Figure 6 left).

**Figure 6.**
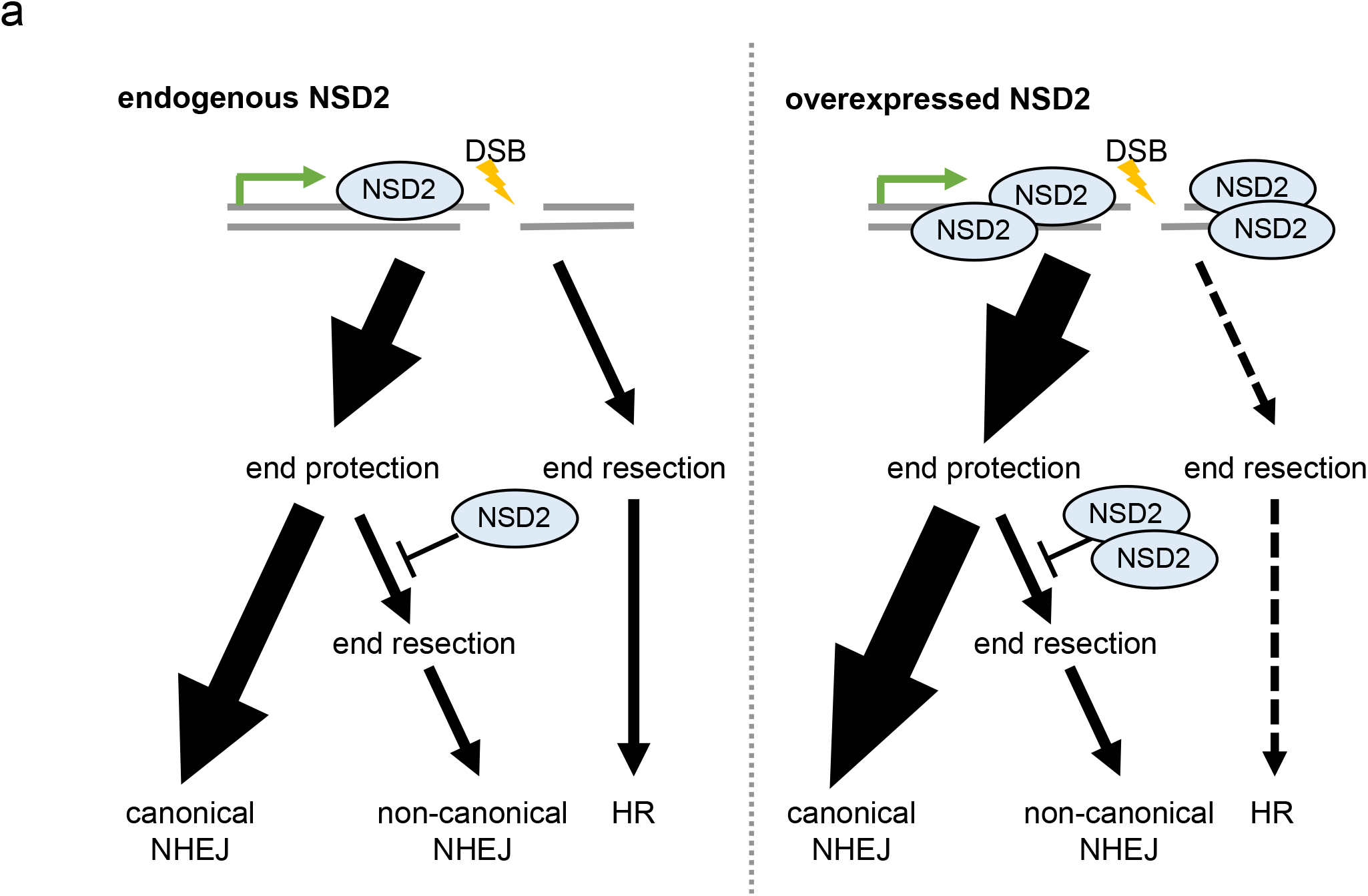
Model for dose-dependent effects of NSD2 on site-specific DSB repair. Endogenous NSD2 localized at actively transcribed regions represses DNA ends resection during NHEJ (left). Excess amounts of NSD2 repress end-resection, promote c-NHEJ and suppress HR (right). DSBs are mainly repaired by NHEJ at the NSD2-bound active genes.

In contrast to endogenous NSD2, our results indicated that the overexpression of NSD2 affected the pathway choice for the repair of DSBs (Figure 6 right). NSD2 overexpression promoted NHEJ (Figure 3d) and suppressed HR (Figure 3b) at the active *TK* locus. Consistent with the results from the DSB reporter experiments, we observed a significant reduction in IR-induced RAD51 foci formation in cells overexpressing NSD2 (Figure 5), which indicates that end resection is being repressed. Indeed, our amplicon-seq analysis showed that NSD2 overexpression reduced the deletion mutant frequency, while we also did not detect any long (>20 bp) deletions around the I-*Sce*I site in these cells (Figure S1). Recent studies have shown an alteration in chromatin domain structures following NSD2 overexpression (Lhoumaud et al., 2019), while NSD2 has enhanced the liquid–liquid phase separations of factors (Liu et al., 2021, Peng et al., 2023). NSD2 dose-dependent effects on the DSB repair pathways might reflect nuclear structure reorganization, which is mediated by an excess amount of NSD2.

The levels of H3K36me2/3, although not H4K20me2, clearly correlated with NSD2 expression levels, thereby indicating that NSD2 is a major H3K36 specific methyltransferase in TK6 cells (Figure 2b, 2d, and 2f). H3K36me is solely catalyzed in yeast by Set2, which maintains chromatin integrity by suppressing histone exchange at active gene regions to repress inappropriate transcriptional initiation (Carrozza et.al., 2005, Venkatesh et al., 2012). Although further studies are required to identify the functional link between the H3K36me activity and control mechanisms in DSB repair by NSD2, our results demonstrate that NSD2 suppresses error-prone non-c-NHEJ, while NSD2 overexpression enhances DSB repair by c-NHEJ. Thus, we propose that NSD2 functions to maintain chromatin integrity during DSB repair.

## 4 Experimental procedures

### 4.1 Cell culture

GM0637, an SV40-transformed human fibroblast cell was cultured in Dulbecco’s modified Eagle’s medium supplemented with 10% fetal calf serum, 100U/mL penicillin, and 100 μg/mL streptomycin. Human lymphoblastoid TK6 cell lines, TSCER2, TSCE105 and TSCE5 (Honma et al., 2007), were cultured in RPMI-1640 medium (Nacalai Tesque) supplemented with 10% fetal bovine serum, 200 μg/mL sodium pyruvate, 100U/mL penicillin, and 100 μg/mL streptomycin.

### 4.2 Plasmids

To express EGFP-NSD2 fusion proteins in GM0637 cells, the mouse *Nsd2* coding region was introduced into pCAGIP-EGFP-gw with Gateway Technology (Invitrogen) (Nimura et al., 2006, 2009).

### 4.3 UVA-microirradiation and immunofluorescence analysis

UVA microirradiation and immunofluorescence analysis was performed as previously described (Shima et al., 2013). The 355 nm line of the UVA-laser was used to introduce DSBs in GM0637 cells. Cells were fixed with 4% paraformaldehyde in PBS for 10 min and then permeabilized in PBS containing 0.5% Triton X-100 for 10 min. The cells were incubated anti-NSD2 antibody (Abcam, ab75359) and anti-53BP1 antibody (Novus, NB100-304) in PBS containing BSA at 37°C for 30 min. Nuclei were stained with 0.5 mg/ml DAPI (Sigma).

### 4.4 Microirradiation iFRAP and FRAP

The iFRAP and FRAP analyses with UVA microirradiation of EGFP-NSD2 were performed as previously described (Ikura, et al., 2007). For the FRAP analysis of EGFP-NSD2 with microirradiation, the 355 nm line of the UV laser was used. The FRAP analyses with the UVA microirradiation for EGFP-NSD2 dynamics were performed as previously described (Ikura, et al., 2007).

### 4.5 Generation of *NSD2*^-/-^ TK6 cells

To generate human *NSD2*^−/−^ TK6 cells, we designed a guide RNA target for CRISPR/Cas9 genome editing in combination with gene targeting constructs. For disruption of *NSD2*, DNA fragments were obtained by PCR from TK6 genomic DNA using the following primers: 5′-ATTCGAGCTCGGTACGGCATCTCTGGTCCTTACTGTAC-3′ and 5′- CCGATCCCATATTGGGCTTCCGTTCTTAGAACACTATGG-3′ for the left arm, 5′- TGCAATAAACAAGTTGCATAAAGATGAAGCAGGCACCAG-3′ and 5′- GCTTGCATGCCTGCAACCTGGCTGAGGAGACAGTAAGT-3′ for the right arm. The neomycin resistance marker gene (*NEO^R^*) was amplified from DT-A-pA/loxP/PGK- *NEO^R^*-pA/loxP vector by PCR using primers 5′-CCAATATGGGATCGGCCATTGA-3′ and 5′-TATCATGTCTGGATCTTCTGCAGACTTACAGCGGATCC-3′. The *simian virus 40 polyadenylation signal* sequence was inserted downstream of the resulting *neo^R^* fragment by overlapping PCR. Then, the amplified fragments of the left arm, right arm, and *neo^R^* were cloned into pUC19L vector by using a GeneArt Seamless cloning kit (Thermo Fisher Scientific). The resulting targeting vector was named pUC-*NSD2-NEO^R^*. In the CRISPR/Cas9 system, the guide RNA (gRNA) sequence for *NSD2* was designed and cloned as double-stranded oligonucleotides (5′-CACCGTTCTAAGAACGGAAGCATC- 3′/5′-AAACGATGCTTCCGTTCTTAGAAC-3′) into the *Bbs*I site of the pX330 vector as described previously (Cong L. et al., 2013). TK6 cells were transfected with 2 μg of the targeting vector pUC-*NSD2-neo^R^* and 6 μg of the pX330-gRNA vector using the NEPA21 electroporator (Nepa Gene Co. Ltd.) following the manufacturer’s instructions. After 48 h, cells were plated in 96-well plates in the presence of G-418 (1 mg/mL). The drug-resistant colonies were picked 10 days after plating and subjected to genomic PCR using the following primers: 5′-CCAATATGGGATCGGCCATTGA-3′ and 5’- AGGAGGCAAAAGACACATCCCT -3’ for a targeted allele with neomycin-resistance cassette or 5′-AGATGAGGTTAAGTGTCCCTGGG-3′ and 5′- AGGAGGCAAAAGACACATCCCT-3′ for identification of homozygous targeting.

### 4.6 Establishment of *NSD2-*overexpressing cells

The *NSD2*-overexpressing cell clone (*NSD2*^-/-^ + *NSD2*) was established by targeted integration of the *NSD2* cDNA into the AAVS1 safe harbor locus. To construct the NSD2 expression plasmid, the coding region of *NSD2* was amplified from the cDNA of HeLa cells by PCR with primers: 5′-GCAGTCGACCTAGTGATGGTGATGATGATG-3’ and 5’-GCCTCTAGATTTGCCCTCTGTGACTCTCCG-3’. Based on the amplified fragment, a construct was designed encoding the Flag-NSD2-IRES-*BSD*^R^ expression cassette under the control of the constitutive EF1α promoter. The resulting plasmid was further digested with *Sph*I and *Bss*HII. The AAVS1 homologous arms were amplified by PCR from TK6 genomic DNA using following primers: 5′- ATTCGAGCTCGGTACCTGAACCTGAGCCAGCTCCCATA-3′ and 5′- GGAGCCTCACGCATGGGACAGATAAAAGTACCCAGAACC-3′ for the left arm, 5′- CCCCGGCCCCGGACGCATCCTTAGGCCTCCTCCTTCCT-3′ and 5′- GCTTGCATGCCTGCAGAAGAGTGAGTTTGCCAAGCAGTC-3′ for the right arm. The DNA fragments of the *NSD2* coding sequence with EF1α promoter and *BSD*^R^ gene, left arm, and right arm were cloned into pUC19L vector by using a GeneArt Seamless cloning kit (Thermo Fisher Scientific). Cells were transfected with 2 μg of the resulting targeting vector pUC-AAVS1-*NSD2-BSD^R^* and 6 μg of the pX330-gRNA-AAVS1 vector containing the guide sequence (5′-CACCGTCACCAATCCTGTCCCTAG-3′/5′- AAACCTAGGGACAGGATTGGTGAC-3′) using the NEPA21 electroporator (Nepa Gene Co. Ltd.). After 48 h, cells were plated in 96-well plates in the presence of blasticidin S hydrochloride (10 μg/mL). The drug-resistant colonies were picked 10 days after plating and subjected to genomic PCR using the following primers: 5′- GACTTCCCCTCTTCCGATGTTGA-3′ and 5’-GGGTTTCAGTGCTAAAACTAGGC-3’ for a targeted allele with blasticidin-resistance cassette.

### 4.7 Western blot analysis

Proteins were separated on a by 12% SDS polyacrylamide electrophoresis and transferred to a PVDF membrane (Millipore). Membranes were incubated in 3% skim milk in wash buffer (2 mM Tris-Cl, pH 8.0, 0.02% NaCl, and 0.05% Tween 20) for 1 h. Membranes were then incubated with the indicated antibodies in blocking buffer for overnight at 4 °C. Membranes were rinsed in wash buffer, and primary antibodies were detected with horseradish peroxidase-conjugated anti-rabbit or anti-mouse (Amersham Biosciences) secondary antibodies. The antibodies used for western blotting are mouse anti-NSD2 (Abcam ab75359), rabbit anti-H3K36me2 (Abcam ab9049) mouse anti-H3K36me3, anti-H3K27-me3, anti-H4K20me2 (gifts from Dr. Kimura) and HRP-linked anti-mouse or rabbit IgG antibody (Cytiva NA9310 or NA9340). Chemiluminescence was detected by LAS 4000 mini (GE Healthcare).

### 4.8 DNA repair assays of I-*Sce*I-induced DSBs

Three TK-6-derived lines, TSCER2, TSCE105 and TSCE5, were used to measure the frequency of HR, NHEJ and alt/-NHEJ events, respectively (Honma2003,2007) as described previously. Briefly, 50 μg I-*Sce*I expression vectors were electroporated into 5 × 10^6^ TK6 cells using Amaxa^TM^ nucleofector (Amaxa Biosystems). After incubation for 72 h at 37°C, cells were seeded into 96-well plates in the presence of hypoxanthin, aminopterin, and thymidine (HAT) for isolating TK-proficient revertants or trifluorothymidine (TFT) for isolating TK-deficient mutants. Drug resistant colonies were counted 4 week later. The revertants and mutant frequencies were calculated as previously reported.

### 4.9 Chromatin immunoprecipitation-quantitative PCR (ChIP-qPCR)

ChIP assays were performed from approximately 5 × 10^6^ cells. Cells were cross-linked with 1% formaldehyde for 10 min at room temperature and formaldehyde was quenched by the addition of 2.5 M glycine to a final concentration of 0.125 M. Cross-linked chromatin was sonicated using a Picoruptor (Diagenode) (10 cycles 30 s ON, 30 s OFF). 2 μg of anti-NSD2 antibody (ab75359) and 20 μl of Dynabeads M-280 sheep anti-mouse IgG (Invitrogen) were mixed in immunoprecipitation dilution buffer and incubated for 6 h at 4 °C. After washing with immunoprecipitation dilution buffer, antibody-binding beads were added to the sonicated chromatin sample and incubated overnight at 4 °C. Beads were washed and chromatin was eluted, followed by reversal of the cross-linking and DNA purification. Chromatin-immunoprecipitated DNA was dissolved in EB buffer (Qiagen). Quantitative PCR was performed with THUNDERBIRD Next SYBR qPCR (TOYOBO) on a CFX384 Touch Real-Time PCR Detection System (BIO RAD). The sequence of the primers are as follows. I-SceI -1kb site Fw: 5′-GTAGAAGGCAGACCCCAGAC-3′, I-SceI-1kb site Rv: 5′-TGAGGTTGGTCAAGAGGTGG-3′, I-sceI +1kb site Fw: 5′-CAGATCCCAGGCCACCAAG-3′, I-SceI +1kb site Rv: 5′- ATGCTCCCTCCTCTCCTACC-3′, I-SceI site Fw: 5′-GTGGACACAGTCCCCCG-3′, I- SceI site Rv: 5′-GATCTGACAACTCCGGCCAT-3′, ARPP21 Fw: 5′- TTGTTTGGGTCTCTGGTAAGCA-3′, ARPP21 Rv: 5′- TGAGGTTACAGGGCAGTCAAGA-3′.

### 4.10 Irradiation and Immunofluorescence analysis

TK6 cells were irradiated with 5 Gy of X-ray using an X-ray generator (CP-160, Faxitron). Cells were attached to the cover slip using cytospin. Nuclei were pre-extracted for 10 min at 4 °C in CSK buffer [10 mM HEPES-KOH (pH 7.4),300 mM sucrose, 50 mM NaCl, 3 mM MgCl_2_, 1 mM EGTA, 0.5% Triton X-100], fixed in 2% paraformaldehyde for 10 min at RT and permeabilized with 0.5% Triton X-100. The nuclei were incubated with anti-RAD51 antibody (BioAcademia 70-002) and anti-γH2AX antibody (Millipore 05-636) for 30 min at 37◦C, and then incubated with Cy3-conjugated secondary antibody and FITC-conjugated secondary antibody for 30 min at 37°C. Coverslips were mounted onto slides using fluorescence mounting medium containing DAPI (Vector Laboratories Inc.). The samples were assessed with an Axio Imager Z2 microscope (Carl Zeiss, Germany), equipped with the Metafer4 software (MetaSystems, Germany). RAD51 and γH2AX foci were automatically counted in at least 1000 nuclei. The number of foci was divided by the number of total nuclei and expressed as foci / nucleus.

## Acknowledgments

We would like to thank Dr. Hiroshi Kimura (Tokyo Institute of Technology, Japan) for providing the antibodies used to identify the modified histones, Dr. Takeshi Yasuda and Dr. Nakako Nakajima (National Institutes for Quantum Science and Technology) for their helpful discussion, Ms. Akiko Ukai (National Institute of Health Sciences) for technical supports, and members of our laboratory for their support and discussion. This work was supported by JSPS KAKENHI Grant Numbers 15H01345 (K.U), 16K16195 (A.S.) and 19K12339 (A.S.), and grants from Takeda Science Foundation, the Program of the Network-type Joint Usage/Research Center for Radiation Disaster Medical Science and Initiative for Realizing Diversity in the Research Environment.

