## Supplementary material for "Dose-dependent effects of histone methyltransferase NSD2 on site-specific double-strand break repair": Figure S1

**Supplementary Figure S1**

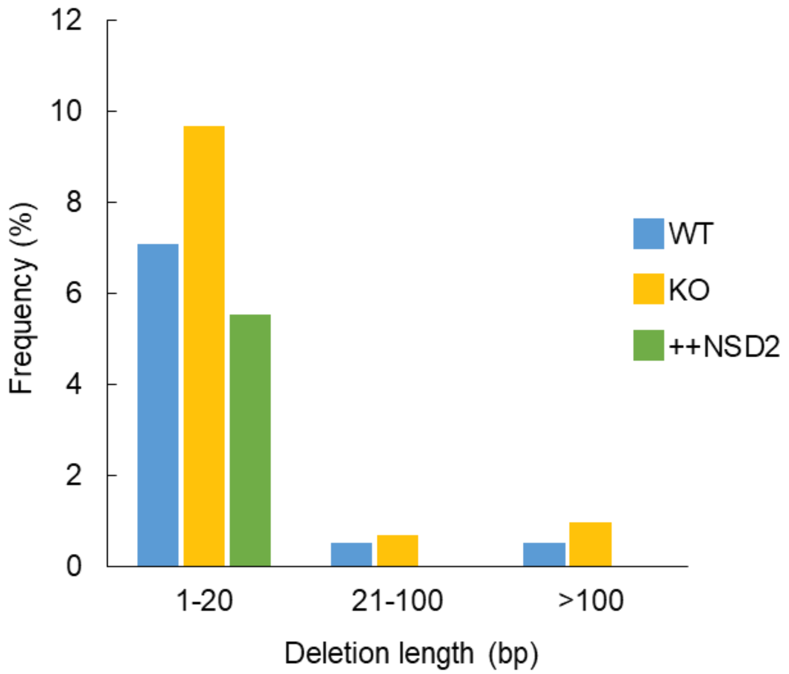


**Supplementary Figure S1. NSD2 dose-dependently represses DNA deletions around the I-SceI site after DSB induction.** Deletion mutation frequencies of wildtype (WT), *NSD2*$^{-/-}$ (KO), and NSD2-overexpressing (++NSD2) TSCE5 cells were analyzed by amplicon-seq. Mutation frequency = (number of reads of a given deletion length) / (total number of reads aligned).

**Supplementary Method**

Amplicon-seq

To isolate I-*Sce*I expressing cells, 48 μg I-*Sce*I expression vectors were co-electroporated with 2 μg GFP expression vectors into 5 x $10^6$ TSCE5 cells using Amaxa™ nucleofector (Amaxa Biosystems). After incubation for 48 h at 37 °C, GFP-positive cells were sorted using a SH800 cell sorter (SONY). Isolated cells were treated with RNase A and proteinase K, and then, the genomic DNA was purified by ethanol precipitation.

The *TK1* intron 4 and exon 5 region fragments, which contain the I-*Sce*I site, were amplified from the genomic DNA of $NSD2^{+/+}$, $NSD2^{-/-}$, and $NSD2^{-/-}$ +NSD2 cells by PCR, using the indicated gene-specific primers. The Illumina adapter and index sequences were attached at the 5' and 3' termini by the additional tailed-PCR with xGen NGS Adapters and Indexing primers (Integrated DNA Technologies, Inc). The constructed libraries were analyzed using D1000 ScreenTape and the Tapestation system (Agilent Technologies, Inc). The 61 bp at the 5' and 3' termini of the constructed libraries were sequenced using NovaSeq 6000 platform (Illumina, Inc). The gene-specific primer sequences used in the experiments are as follows: 5'-TCG TCG GCA GCG TCA GAT GTG TAT AAG AGA CAG CAA ATG GCC GGA GTT GTC-3' and 5'-GTC TCG TGG GCT CGG AGA TGT GTA TAA GAG ACA GCA GAG CTG GTG TCT CTG TGC-3'.

### Amplicon-seq analysis

For each Amplicon-seq paired read, the 19th base from the 5’-end of the I-*Sce*I cut site (5’-GATCCATTACCCTGTTATCCCTACTCTCGAG-3’) was defined as the +1 position and the intact amplicon contained a sequence of -37 to +268. The low-quality sequenced read pairs were filtered out using the criteria of more than two base mismatches over the positions ranging from either -19 to -11 or +212 to +248. The deletion length was manually calculated by aligning 9 successive bases from the 47th base, with counting beginning from the 5'-end of the amplicon via the +10 to +211 sequence of the intact genome.
